# Bone Morphology is Regulated Modularly by Global and Regional Genetic Programs

**DOI:** 10.1101/324293

**Authors:** Shai Eyal, Shiri Kult, Sarah Rubin, Sharon Krief, Kyriel M. Pineault, Deneen M. Wellik, Elazar Zelzer

## Abstract

During skeletogenesis, a variety of protrusions of different shapes and sizes develop on the surfaces of long bones. These superstructures provide stable anchoring sites for ligaments and tendons during the assembly of the musculoskeletal system. Despite their importance, the mechanism by which superstructures are patterned and ultimately give rise to the unique morphology of each long bone is far from understood. In this work, we provide further evidence that long bones form modularly from *Sox9*^+^ cells, which contribute to their substructure, and from *Sox9*^+^/*Scx*^+^ progenitors that give rise to superstructures. Moreover, we identify components of the genetic program that controls the patterning of *Sox9*^+^/*Scx*^+^ progenitors and show that this program includes both global and regional regulatory modules.

Using light sheet fluorescence microscopy combined with genetic lineage labeling, we mapped the broad contribution of the *Sox9*^+^/*Scx*^+^ progenitors to the formation of bone superstructures. Additionally, by combining literature-based evidence and comparative transcriptomic analysis of different *Sox9*^+^/*Scx*^+^ progenitor populations, we identified genes potentially involved in patterning of bone superstructures. We present evidence indicating that *Gli3* is a global regulator of superstructure patterning, whereas *Pbx1, Pbx2, Hoxa11* and *Hoxd11* act as proximal and distal regulators, respectively. Moreover, by demonstrating a dose-dependent pattern regulation in *Gli3* and *Pbx1* compound mutations, we show that the global and regional regulatory modules work coordinately. Collectively, our results provide strong evidence for genetic regulation of superstructure patterning that further supports the notion that long bone development is a modular process.

## INTRODUCTION

The vertebrate skeleton is composed of numerous bones, each displaying a unique shape and size. Yet, despite this morphological diversity, most bones are formed by a common developmental process, namely endochondral ossification (Berendsen and Olsen, 2015; Cervantes-Diaz et al., 2017; Long and Ornitz, 2013; Olsen et al., 2000). During this process, mesenchymal cells derived from the lateral plate mesoderm under the regulation of transcription factor SRY-box 9 (*Sox9)* condense and differentiate into chondroprogenitors, forming cartilaginous anlagen of the future bones (Kawakami et al., 2006; Zhao et al., 1997). Next, a cascade of chondrocyte differentiation steps will give rise to growth plates on both proximal and distal ends of the anlage. Subsequently, blood vessels invade the cartilage anlage introducing bone-building cells, termed osteoblasts. These cells deposit calcium and ossify the cartilaginous template from the mid-shaft, pursuing the progression of the growth plates (Kronenberg, 2003).

Endochondral ossification has been studied extensively for more than two centuries, generating a vast amount of information on the generic regulation of this process. Nevertheless, the mechanism that grants each bone with its distinctive shape is missing. A hallmark of the unique morphology of each long bone is the superstructures that protrude from its surface known as bone eminences, such as the greater and deltoid tuberosities, greater and lesser trochanters, etc. and condyles, such as the distal lateral and medial epicondyles of the humerus. One of the main functions of bone superstructures is to provide an attachment point for tendons and ligaments, which transmit force from the contracting muscles to the skeleton (Lessa et al., 2008; McHenry and Corruccini, 1975; Polly, 2007).

Interestingly, it was previously shown that the site where tendon attaches to bone is formed by a unique set of progenitors that co-express *Sox9* and scleraxis (*Scx*) (Blitz et al., 2013; Sugimoto et al., 2013). Moreover, it was demonstrated that these cells that contribute to the bony side of the attachment do not descend from the cells that create the bone shaft anlage and are specified after them. Finally, it was demonstrated that these unique progenitors are specifically regulated by the TGFβ and BMP signaling pathways (Blitz et al., 2009; Blitz et al., 2013). In addition to providing a mechanism for the development of tendon-to-bone attachment site, these findings also offer an alternative, modular model for long bone development. According to this model, one set of *Sox9*^+^ cells forms the cylindrical anlage of the future bone shaft, which will serve as the bone substructure, whereas a second pool of *Sox9*^+^/*Scx*^+^ cells will be added onto this substructure to give rise to the different superstructures.

Current literature provides ample evidence in support of the modular model of long bone development, which is based on lineage tracing, temporal initiation and cell differentiation of the *Sox9*^+^/*Scx*^+^ progenitor cells. However, this model is still missing the mechanism that regulates the early patterning of the *Sox9*^+^/*Scx*^+^ progenitor cells, such that superstructures will form on the developing long bone at the right location and size. In this study, we examine these regulatory mechanisms underlying this aspect of the modularity in skeletal development. Three-dimensional (3D) reconstruction of different long bones and their protruding superstructures allowed us to demonstrate modularity in long bone development. Comparative transcriptomic analysis and loss-of-function assays showed that GLI-Kruppel family member GLI3 (*Gli3*) globally regulates the spatial organization of the *Sox9*^+^/*Scx*^+^ progenitors in both forelimb and hindlimb, whereas homeobox a11 (*Hoxa11*) and *Hoxd11* regulates patterning of distal superstructures and pre B cell leukemia homeobox 1 (*Pbx1*) and *Pbx2* regulate proximal superstructure progenitors. Overall, we provide cellular evidence for the existence of a patterning mechanism that involves both global and regional regulation and highlight some of the genes that facilitate the patterning of superstructures along the developing long bones.

## RESULTS

### *Sox9*^+^/*Scx*^+^ superstructure progenitors contribute extensively to the morphology of long bone anlagen

To better understand the scope of modularity in long bone morphogenesis, we performed pulse-chase cell lineage experiments by crossing either *Sox9-CreER*^*T2*^ or collagen type II alpha 1 (*Col2a1*)-*CreER*^*T2*^ transgenic reporter mice with *Rosa26-tdTomato* reporter mice (Madisen et al., 2010; Nakamura et al., 2006; Soeda et al., 2010). Pregnant females were administered a single dose of 0.03 mg/gr tamoxifen/body weight at either E10.5 or E11.5. Previously, we showed that this time window allows the exclusive labeling of cells of the cylindrical bone substructure, leaving the superstructure cells unlabeled (described in detail in (Blitz et al., 2013; Eyal et al., 2015)). To map comprehensively the unlabeled superstructure cells in different skeletal elements, we have established a 3D imaging pipeline that includes clearing of the labeled limbs optically, light-sheet fluorescence microscopy and, finally, reconstruction of obtained images into a 3D object (Treweek et al., 2015; Yang et al., 2014). To better visualize the superstructures, the whole-mount limbs were immunostained for collagen type II alpha 1 (COL2A1).

As seen in Figure 1A,B (and Supplementary Movies M1-M3), in E14.5 *Sox9-tdTomato*^+^ or E15.5 *Col2a1-tdTomato*^+^ limbs, all superstructures of different long bones were *tdTomato*-negative, including greater, lesser and deltoid tuberosities, olecranon, and greater, lesser and third trochanters. Interestingly, in addition to bone eminences, we observed various condyles and sesamoid bones that were also *tdTomato*-negative, including distal medial and lateral humeral epicondyles, medial tibial condyle, patella, and medial and lateral fabella. These results demonstrate the generality of the modular process of long bone morphogenesis in the limb.

**Figure 1.**
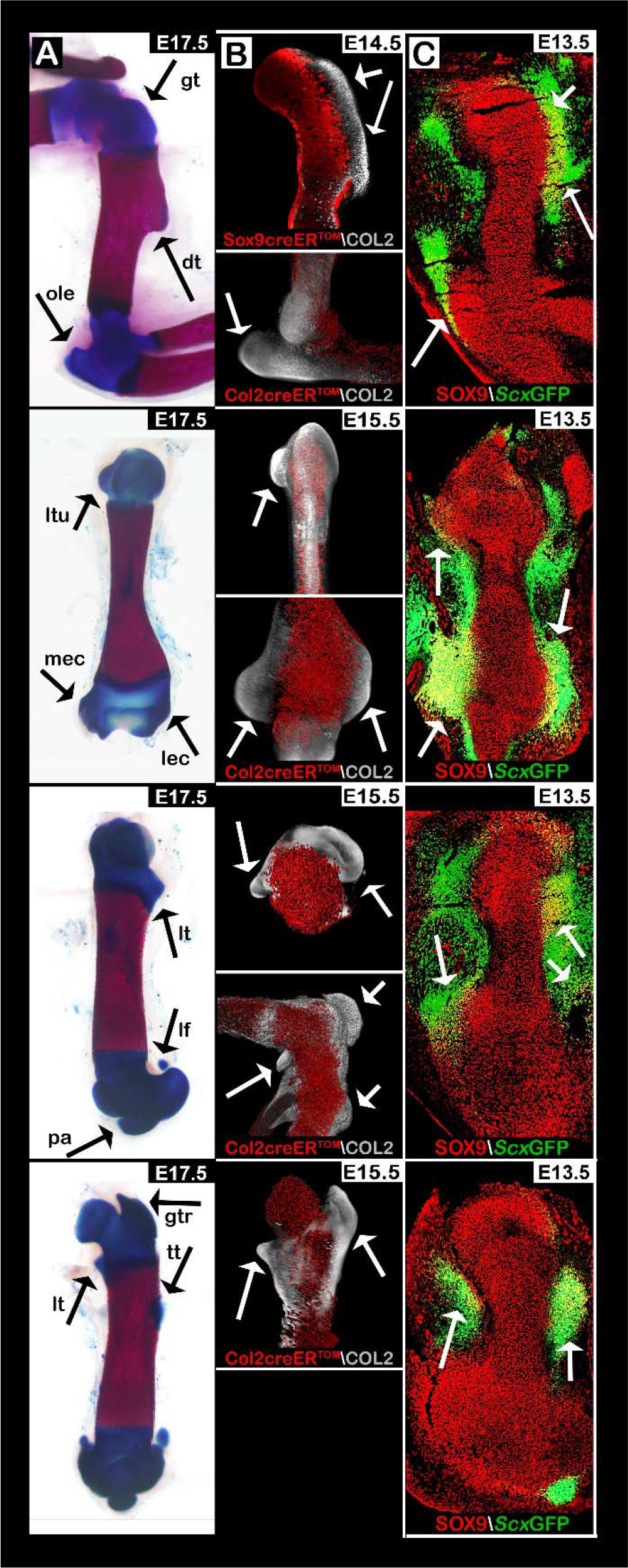
Long bone morphology is affected extensively by numerous pools of *Sox9*^+^/*Scx*^+^ progenitors. Images showing sagittal and coronal views of the superstructures that form along the forelimb and hindlimb long bones. **(A)** Skeletal preparation of E17.5 WT limbs. Superstructures are indicated by black arrows. **(B)** Maximum intensity projection (MIP) images of E14.5 forelimb from *Sox9-Cre*^*ER*^-*tdTomato* transgenic embryos and E15.5 fore- and hindlimbs from *Col2a1-Cre*^*ER*^-*tdTomato* transgenic embryos. Whole-mount limbs were stained against COL2A1, imaged using light sheet microscopy and reconstructed. Whereas first-wave *Sox9*^+^ progenitors were labeled by tdTomato, the secondary wave of *Sox9*^+^/*Scx*^+^ progenitors remained unstained. tdTomato^−^ precursors contributed exclusively to superstructure formation (white arrows). **(C)** Sagittal and coronal sections through E13.5 fore- and hindlimbs from *ScxGFP* transgenic embryos stained against SOX9. *Sox9* and *Scx* coexpressing cells are indicated by white arrows. Abbreviations: gt, greater tuberosity; dt, deltoid tuberosity; ole, olecranon; ltu, lesser tuberosity; mec, medial epicondyle; lec, lateral epicondyle; lt, lesser trochanter; lf, lateral fabella; pa, patella; gtr, greater trochanter; tt, third trochanter.

To further demonstrate this modularity, we examined sagittal and coronal sections of E13.5 fore- and hindlimbs from *ScxGFP* transgenic embryos (Pryce et al., 2007) that were stained using antibodies against SOX9. As seen in Figure 1C, we observed an abundance of *Sox9*^+^/*Scx*^+^ progenitors at presumable anatomical locations of various superstructures along the shafts of different bones. Taken together, these results indicate the extensive contribution of *Sox9*^+^/*Scx*^+^ superstructure progenitors to the 3D morphology of cartilaginous anlagen, highlighting the modularity of this process both temporally and lineage-wise.

### *Gli3* is a global regulator of *Sox9*^+^/*Scx*^+^ progenitors patterning during superstructure development

An immediate implication of the modular model is the existence of a mechanism that regulates the patterning of the *Sox9*^+^/*Scx*^+^ progenitors to position the superstructure correctly along the bone substructure. To uncover this mechanism, we first searched the literature for mutations leading to observable superstructure patterning abnormalities. We found that the skeletons of *Gli3*-null embryos, previously known as extra toes (*Xt*), exhibit a variety of superstructure abnormalities (Johnson, 1967). To characterize these defects, we crossed *Gli3* heterozygous mice and examined skeletal preparations from E17.5 embryos. As seen in Figure 2A-I, both fore- and hindlimb of *Gli3*^*null*^ embryos exhibited numerous dysplastic superstructures, including lesser and deltoid tuberosity, olecranon, patella, and additional ectopic sesamoids. Notably, the aberrant patterning of the deltoid tuberosity ranged in severity from mildly abnormal morphology to complete separation from the humeral shaft (compare left and right forelimbs in Fig. 2F and G, respectively). We also examined *Prx1-Cre*;*Gli3*^*fl/fl*^ embryos, in which *Gli3* was conditionally knocked out (cKO) in the limbs (Blaess et al., 2008; Logan et al., 2002). Similarly to *Gli3*^*null*^ mutants, limb-specific *Gli3* cKO also resulted in diverse abnormally patterned superstructures, such as induction of supernumerary sesamoids around the knee, dysplastic medial tibial condyle and dysplastic ulnar olecranon and coronoid processes (Fig. 2J-O).

**Figure 2.**
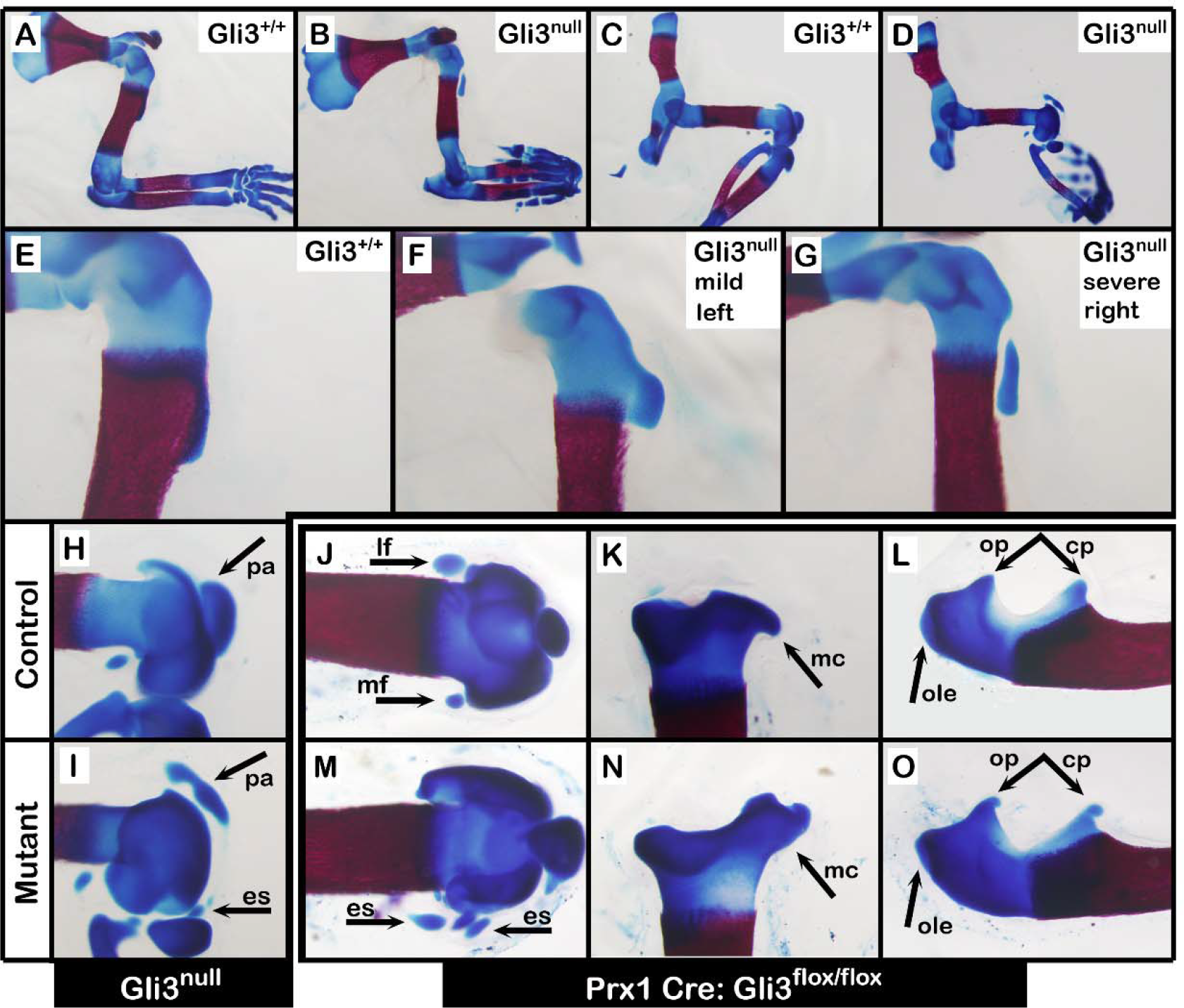
*Gli3* regulates the patterning of superstructures globally. **(A-O)** Skeletal preparations of E17.5 fore- and hindlimbs from *Gli3* KO and control embryos (A-I) and from limb-specific *Prx1-Cre;Gli3*^*floxed*^ cKO and control embryos (J-O). Affected superstructures are indicated by black arrows. Abbreviations: pa, patella; es, ectopic sesamoids; ole, olecranon; op, olecranon process; cp, coronoid process; mc, medial tibial condyle; lf, lateral fabella; mf, medial fabella.

To study the possibility that *Gli3* regulate the patterning of superstructure progenitors, we sought to examine the distribution of *Sox9*^+^/*Scx*^+^ cells in *Gli3*^*null*^ limbs. For that, we crossed heterozygous *Gli3* mice with *Gli3*^+/-^*; ScxGFP* transgenic reporter mice and stained sagittal sections of E13.5 fore- and hindlimbs against SOX9 to highlight chondrocytes. As seen in Figure 3A-F, the spatial distribution of *Sox9*^+^/*Scx*^+^ progenitors in *Gli3*^*null*^ limbs was abnormal. Whereas in control limbs these progenitors were patterned in juxtaposition to the humeral shaft (Fig. 3A; dotted line), in the mutant they were laterally spread and less condensed (Fig. 3D; dotted line). To understand how the abnormal distribution of *Sox9*^+^/*Scx*^+^ progenitors is translated into superstructures abnormalities, we examined the spatiotemporal differentiation of deltoid tuberosity precursors in E13.5 and E16.5 sagittal sections from control and *Gli3* KO forelimbs. Sections were stained using antibodies against SOX9 and COL2A1, which comprises the collagen-rich matrix deposited by differentiated chondrocytes (Fig. 3G-L). As expected, at E13.5 we repeatedly observed the lateral spreading of deltoid tuberosity precursors (Fig. 3H and I; dotted lines). Moreover, we noticed that the degree of cellular lateralization varied both in pattern and in distance from the humeral shaft (Fig. 3H and I; white bars). At E16.5, the abnormal deltoid tuberosity morphologies were clearly evident and ranged from dysplastic to separate deltoid tuberosity (Fig. 3K, L, respectively). We concluded that *Gli3* is necessary for correct spatial organization of deltoid tuberosity *Sox9*^+^/*Scx*^+^ precursors and, thus, for superstructure patterning.

**Figure 3.**
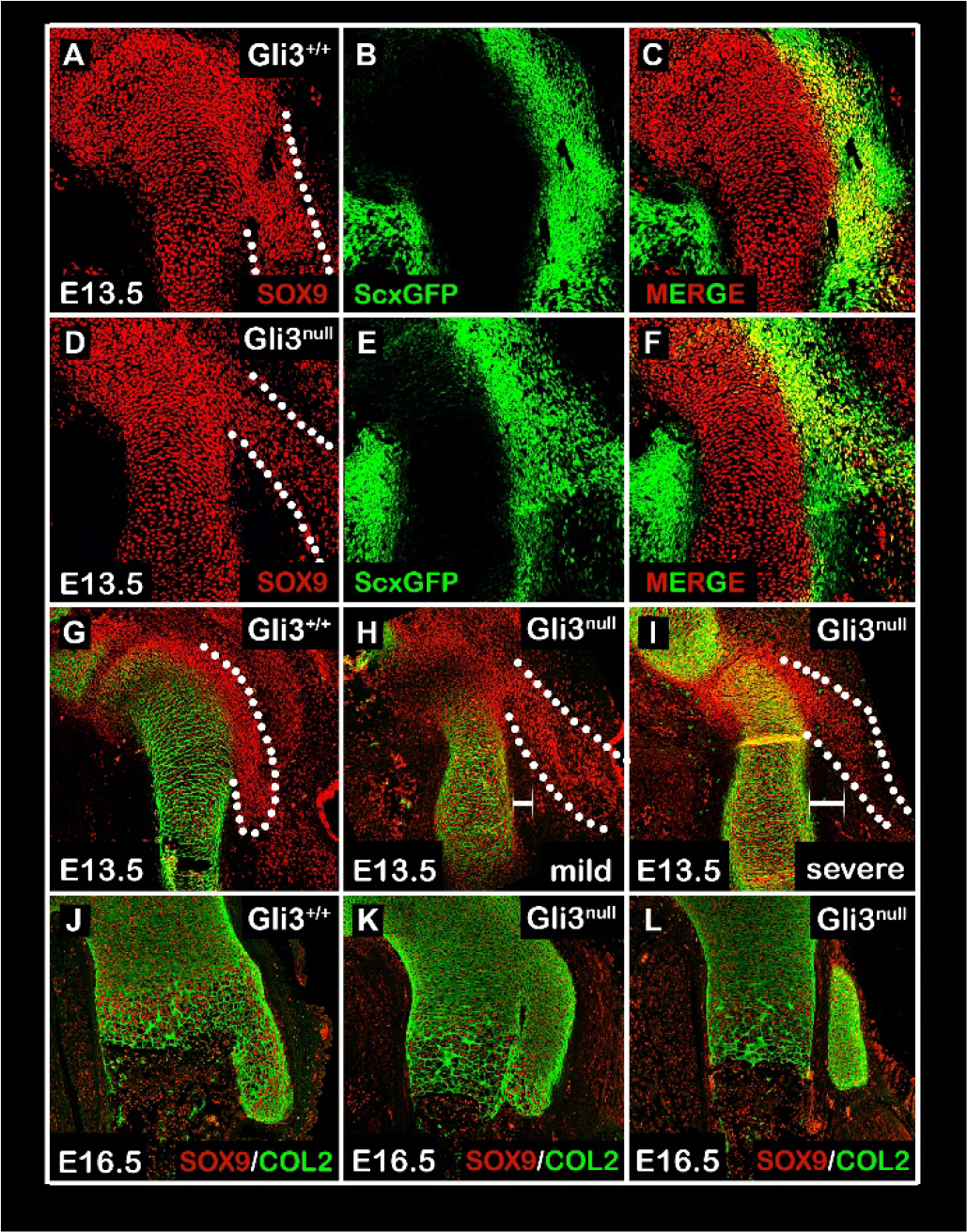
Organization but not specification of *Sox9*+/*Scx*+ progenitors is regulated by *Gli3.* **(A-F)** Sagittal sections through the proximal humeri of E13.5 *Gli3*;*ScxGFP* transgenic embryos that were stained against SOX9. Whereas *Sox9* and *Scx* co-expressing cells are observed in control and mutant embryos (C and F), their spatial organization is abnormal in *Gli3*^*null*^ mutants (A and D, dotted lines). **(G-L)** Sagittal sections through the proximal humeri of E13.5 (G-I) and E16.5 (J-L) *Gli3* KO and control embryos that were stained against SOX9 and COL2A1. At E13.5, spatial organization of DT precursors is abnormal in mutants (H and I; dotted lines), displaying varying degrees of lateralization and distance from the humeral shaft (H and I; white bars). In E16.5 mutants, aberrant DT morphology ranges from aplasia (K) to detachment from the humeral shaft (L).

### Comparative transcriptomic analysis of proximal and distal *Sox9*^+^/*Scx*^+^ progenitors predicts regional regulation of superstructure patterning

Although *Gli3* had a global effect on superstructure patterning, we suspected that regional regulatory circuitry also exists, such that patterning of specific superstructures could be regulated independently. To search for genes potentially involved in this mechanism, we developed a protocol for prospective isolation of *Sox9-tdTomato*^+^/*ScxGFP*^+^ progenitors by FACS. Moreover, to enable distinction between global and regional patterning regulators, we designed our system to allow separate isolation of proximal and distal cell populations. To this end, we crossed *Sox9-CreER*^*T2*^*;Rosa26-tdTomato* reporter mice with *ScxGFP* transgenic reporter mice. Pregnant females were administered a single dose of 0.03 mg/gr tamoxifen/body weight at E12.0 (Fig. 4A). Next, we harvested E13.5 *Sox9-tdTomato*^+^/*ScxGFP*^+^ embryos and micro-dissected their forelimbs at the shoulder level, above the presumable DT, and at flanking sides of the elbow (Fig. 4B i and ii). Finally, the dissected segments were collected into individual tubes, homogenized and then prospectively isolated by FACS (Fig. 4Biii). For each proximal and distal segment, we collected three biological repeats from three separate litters. Each sample contained 7-9% *Sox9-tdTomato*^+^/*ScxGFP*^+^ progenitors of total cell population, of which 10,000 cells were collected for sequencing (Fig. 4C). Further information regarding controls, gate settings and instrument configuration is found in Fig. 4D and in Materials and Methods.

**Figure 4.**
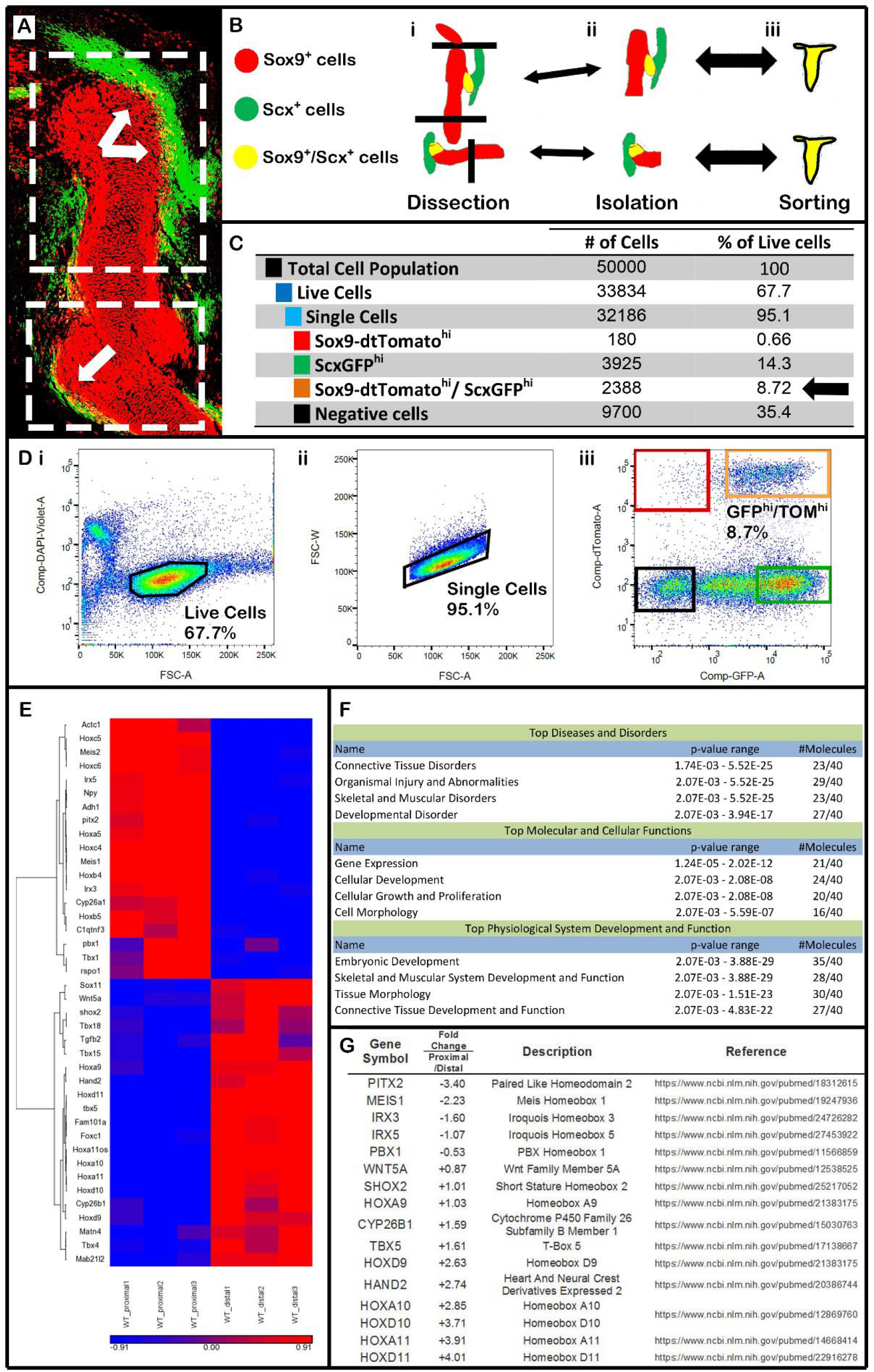
Comparative transcriptomic analysis highlights regional regulation of super-structure patterning. **(A)** Sagittal section through the forelimb of E13.5 *Sox9-CreER*^*tdTomato*^*;Scx-GFP* transgenic reporter embryo following tamoxifen administration at E12.0. *Sox9* and *Scx* co-expressing cells are indicated by white arrows. Dashed white rectangles indicate the proximal and distal regions of interest that were dissected for FACS and transcriptome analysis. **(B)** Schematic diagram of sample preparation. Labeled forelimbs were dissected (i), the proximal and distal segment cells were isolated (ii) and sorted independently (iii). **(C)** A table summarizing cell type distribution among 50,000 collected and FACS-sorted cells. The number and percentage of *Sox9-tdTomato*^+^/*ScxGFP*^+^ progenitors are indicated by a black arrow. **(D)** Illustrations of control and gating settings. (i) Live cells were controlled for by DAPI staining. (ii) Single cells were gated according to droplet area (FSC-A) vs. width (FSC-W). (iii) Gating settings for *Sox9-tdTomato*^+^/*ScxGFP*^+^ progenitors resulted in collection of between 7-9% of total living single-cell population (orange rectangle). (E) Heat map showing the clustering of 40 of the most differentially expressed genes between proximal and distal *Sox9-tdTomato*^+^/*ScxGFP*^+^ progenitors. **(F)** Using Ingenuity software, these genes were annotated and found to be biologically relevant to limb development and tissue morphology. **(G)** Sixteen of these genes have previously been implicated in regional regulation of superstructure patterning.

Comparison between the transcriptomes of proximal and distal *Sox9-tdTomato*^+^/*ScxGFP*^+^; progenitors revealed 561 differentially expressed genes with at least two-fold change in expression levels between segments (Supplementary Table T1). We further analyzed 40 of these genes that were considered most likely to be involved, based on the difference in expression levels, relevant biological function and annotation as developmental genes (Fig. 4E and 4F). Notably, 16 of these 40 candidate genes (referenced in Fig. 4G) have previously been implicated in superstructure development, providing a strong validation for the effectiveness of our strategy.

### *Hoxa11*, *Hoxd11* and *Pbx1* genes regionally regulate distal or proximal superstructure patterning

Our transcriptome analysis revealed that *Hoxa11* and *Hoxd11* were highly expressed in distal *Sox9-tdTomato*^+^/*ScxGFP*^+^ progenitors and scarcely expressed in proximal cells. Intriguingly, a previous study has demonstrated that *Hoxa11/Hoxd11* compound mutation (*Hox11*^*aadd*^) resulted in diverse distal forelimb abnormalities, such as the formation of a detached olecranon resembling a sesamoid bone, whereas the proximal limb segment, including the DT, was not affected (Koyama et al., 2010). Nevertheless, the mechanism underlying the abnormal olecranon development remained to be elucidated. Therefore, we proceeded to study this process in *Hox11*^*aadd*^ mutant embryos.

To that end, we crossed double-heterozygous *Hox11*^*aaDd*^ mice and harvested embryos at E13.5 and E17.5 (Wellik and Capecchi, 2003). Examination of skeletal preparations from E17.5 control and mutant embryos validated the abnormal development of the olecranon (Fig. 5A,A’). Next, we validated the differential expression of *Hoxd11* by fluorescent in situ hybridization (FISH) assay on sagittal sections of E13.5 forelimbs. In agreement with our transcriptome analysis results, *Hoxd11* was highly expressed distally in proximity to the elbow, but scarcely expressed in proximal parts of the limb (Fig. 5B). Furthermore, *Hoxd11* expression was down-regulated in *Hox11*^*aadd*^ mutants (Fig. 5B’). The differential expression of *Hoxa11* at this stage was previously demonstrated (Swinehart et al., 2013). To study the regulatory roles of *Hoxa11* and *Hoxd11* in distal patterning of superstructure progenitors, we examined the patterning of the olecranon in control and mutant embryos. For that, we crossed double-heterozygous *Hox11*^*aaDd*^ mice and stained forelimb sections from E13.5 double-homozygous *Hox11*^*aadd*^ embryos using antibodies against SOX9 and COL2A1, such that the substructure cells were expected to be *Sox9*^+^/*Col2a1*^+^ and the undifferentiated superstructure progenitors to be *Sox9*^+^/*Col2a1*^−^. Results showed that the olecranon progenitors failed to organize in the typical pattern observed in control limbs (Fig. 5C-E’). Specifically, the cells were scattered in a proximolateral direction away from the ulnar shaft (Fig. 5C,C’; dotted lines). Sections from E17.5 *Hox11*^*aadd*^ mutants illustrated that the olecranon precursors differentiated at a distance from the ulnar shaft, creating a distinct skeletal element resembling a sesamoid bone (Fig. 5F,F’). These results suggest that *Hoxa11* and *Hoxd11* are necessary for correct localization of the olecranon progenitors and, thus, for their patterning. Moreover, the proper patterning of superstructure progenitors on the proximal side supports our hypothesis that *Hoxa11* and *Hoxd11* act as regional distal regulators (Sup Fig. 1A-A’).

**Figure 5.**
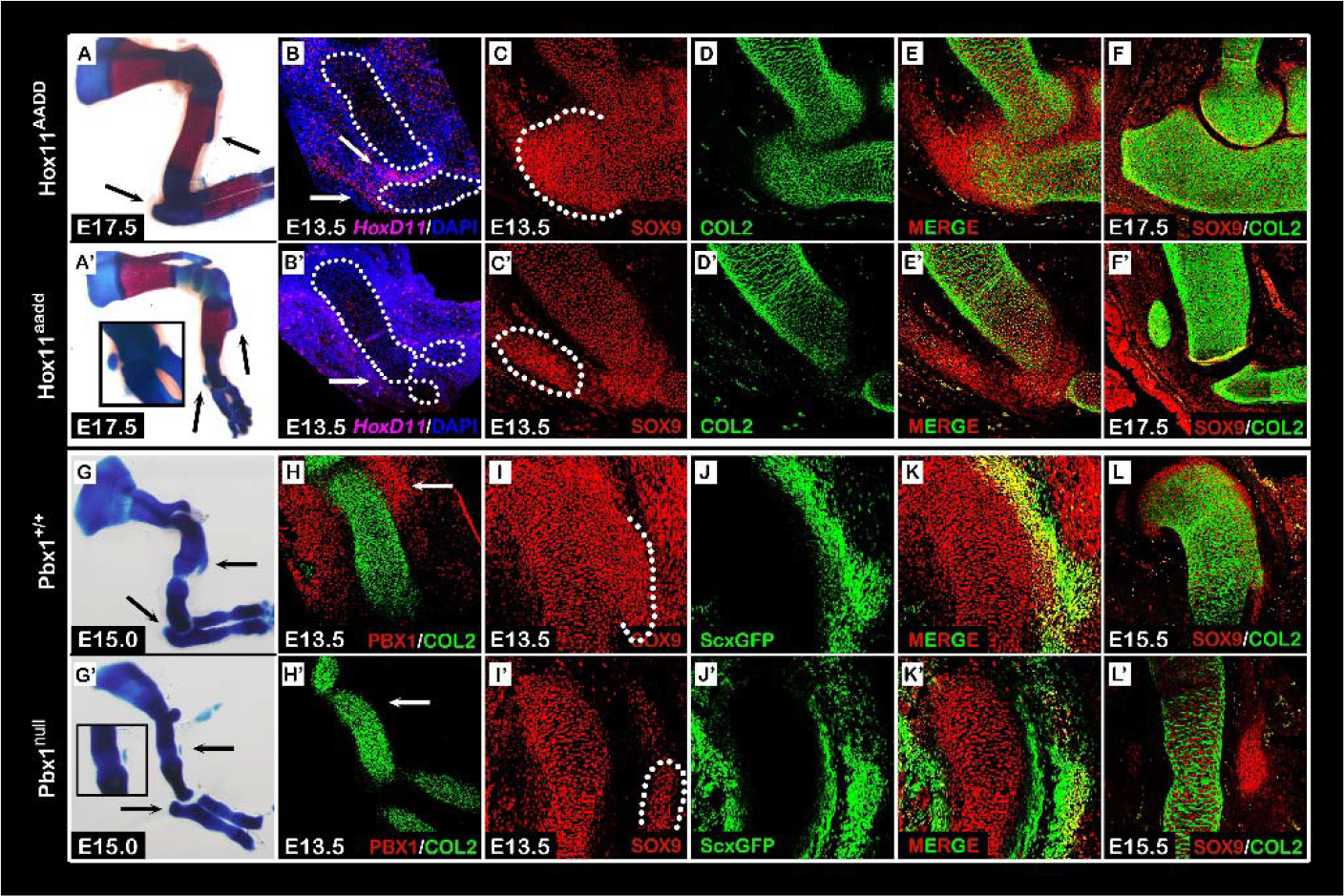
Distal and proximal superstructure patterning is regulated by *HoxA11*, *HoxD11* and *Pbx1.* **(A-F’)** *HoxA11* and *HoxD11* regionally regulate patterning of the distal olecranon. (A,A’): Skeletal preparations of E17.5 forelimbs from compound *HoxA11* and *HoxD11* mutant and control embryos. Distal olecranon but not proximal DT developed abnormally in *Hox11*^*aadd*^ mutant forelimbs (black arrows and enlarged rectangle). (B,B’): Fluorescent in situ hybridization for *Hoxd11* in forelimb sections shows that at E13.5, *Hoxd11* is distally expressed in the elbow region (B, white arrows). Expression of *HoxD11* was downregulated in *Hox11*^*aadd*^ mutant forelimbs (B’). Long bones are demarcated by dotted lines. (C-F’): Sagittal sections through the elbow of E13.5 (C-E’) and E17.5 (F,F’) *Hox11*^*aadd*^ compound mutant and control embryos that were stained against SOX9 and COL2A1. At E13.5, spatial organization of olecranon precursors is abnormal (C and C’; white arrowheads). By E17.5, the developing olecranon of mutant embryos has detached from the ulnar shaft (F,F’). **(G-L’)** *Pbx1* regionally regulates patterning of the proximal DT. (G,G’): Skeletal preparations of E15.0 forelimbs from *Pbx1*-null and control embryos. Proximal DT but not distal olecranon developed abnormally in *Pbx1* mutant forelimbs (black arrows). (H,H’): Staining against PBX1 and COL2A1 in forelimb sections shows that at E13.5, *Pbx1* is proximally expressed at the shoulder region (H, white arrows). Expression of *Pbx1* was downregulated in *Pbx1*^*null*^ mutant forelimbs (H’). (I-K’): Sagittal sections through the proximal humeri of E13.5 *Pbx1*;*ScxGFP* transgenic embryos that were stained against SOX9. Although *Sox9* and *Scx* co-expressing cells are observed in control and mutant embryos (K and K’), their spatial organization was abnormal in *Pbx1*^*null*^ mutants (I and I’, white arrowheads). (L,L’): Sagittal sections through the proximal humeri of E15.5 *Pbx1* mutant and control embryos that were stained against SOX9 and COL2A1. At E15.5, the DT of mutant embryos is detached from the humeral shaft (L,L’).

Another promising candidate gene from our transcriptome analysis was *Pbx1*. In contrast to *Hoxa11* and *Hoxd11, Pbx1* was highly expressed in proximal *Sox9-tdTomato*^+^/*ScxGFP*^+^ progenitors as compared to distal cells. Indeed, *Pbx1*^*null*^ mutation has been shown to affect only proximal elements, such as the DT, whereas the distal limb segment, including the olecranon, was not affected (Selleri et al., 2001). We therefore performed the same analysis on *Pbx1*^*null*^;*ScxGFP* mutants. After validation of the deltoid tuberosity phenotype in skeletal preparations of E15.0 control and mutant embryos (Fig. 5G,G’), we analyzed *Pbx1* expression by immunostaining sagittal sections of E13.5 forelimbs against PBX1 and COL2A1. In agreement with the transcriptome analysis results, *Pbx1* was highly expressed proximally in proximity to the deltoid tuberosity and was less prominent in the distal forelimb (Fig. 5H). As expected, *Pbx1* expression was completely ablated in *Pbx1*^*null*^ mutants (Fig. 5H’).

Next, forelimb sections from *Pbx1*^*null*^;*ScxGFP*^+^ E13.5 embryos were stained using antibodies against SOX9. As seen in Figure 5I-K, the *Sox9*^+^/*Scx*^+^ deltoid tuberosity precursors were specified in both control and mutant limbs; yet, in the mutant they were organized abnormally and scattered laterally. Finally, examination of sections immunostained against SOX9 and COL2A1 at E15.5 showed that in *Pbx1*^*null*^ mutants, the deltoid tuberosity precursors differentiated into a distinct sesamoid-like cartilaginous element parallel to the humeral shaft (Fig. 5L,L’). These results indicate that *Pbx1* is necessary for correct localization of the *Sox9*^+^/*Scx*^+^ deltoid tuberosity precursors and, thus, for their patterning. Importantly, the patterning of superstructure progenitors at the distal region was independent of *Pbx1* regulation (Sup Fig. 1B-B’).

Together, these results indicate that superstructure patterning is regulated regionally through early modulation of the spatial organization of *Sox9*^+^/*Scx*^+^ progenitors by *Hoxa11, Hoxd11* and *Pbx1*.

### Both *Pbx1* and *Pbx2* are involved in patterning of proximal superstructures

*Pbx2* is a paralog of *Pbx1* that is expressed throughout the limb mesenchyme. Previously, it was shown that *Pbx1* and *Pbx2* act in a dosage-dependent manner during proximal limb development (Capellini, 2006). This led us to examine if *Pbx2* co-regulates proximal superstructures patterning with *Pbx1*. To avoid early embryonic lethality, we ablated one allele of *Pbx2* on the background of a limb-specific knockout of *Pbx1* (*Prx1-Cre;Pbx1*^*fl/fl*^*;Pbx2*^+/−^) and compared the phenotype to both WT (*Prx1-Cre*^−^*;Pbx1*^+/+^*;Pbx2*^+/+^) or *Pbx1* cKO (*Prx1-Cre;Pbx1*^*fl/fl*^) embryos (Ficara et al., 2008; Selleri et al., 2004). To this end, sections from E13.5, E15.5 and E16.5 control and mutant forelimbs were immunostained using antibodies against SOX9 and COL2A1 (Fig. 6A-C”). At E13.5, we observed abnormal patterning of deltoid tuberosity precursors in both *Prx1-Cre;Pbx1*^*fl/fl*^ and *Prx1-Cre;Pbx*^*fl/fl*^*;Pbx2*^+/−^ mutant embryos. The precursors were laterally scattered (Fig. 6A’ and A”), comparable to the deltoid tuberosity precursors of *Pbx1*^*null*^ limbs (Fig. 5I’). By E15.5, the deltoid tuberosity precursors of *Prx1-Cre*;*Pbx1*^*fl/fl*^ embryos have fully differentiated, whereas in *Prx1-Cre;Pbx1*^*fl/fl*^*;Pbx2*^+/−^ we observed delayed differentiation (Fig. 6B-B”). Notably, in *Prx1-Cre;Pbx1*^*fl/fl*^ embryos the humerus-DT boundary was populated by *Sox9*^+^ cells that had not been detected in *Prx1-Cre;Pbx1*^*fl/fl*^*;Pbx2*^+/−^ embryos. Moreover, the gap between the humeral shaft and deltoid tuberosity precursors was roughly twice as large in *Prx1-Cre*;*Pbx1*^*fl/fl*^;*Pbx2*^+/−^ limbs than in *Prx1-Cre*;*Pbx1*^*fl/fl*^ limbs. Finally, by E16.5, the deltoid tuberosity of *Prx1-Cre*;*Pbx1*^*fl/fl*^ embryos has attached to the humeral shaft, while the deltoid tuberosity of *Prx1-Cre;Pbx1*^*fl/fl*^*;Pbx2*^+/−^ mutants remained separated (Fig. 6C-C”), as in *Pbx1*^*null*^ mutants (Fig. 5L’). These diverging morphologies were further validated by skeletal preparation made of limbs taken from each mutants (Fig 6. D-D”). Taken together, these results suggest that PBX genes act together in regulating the patterning of proximal superstructures, possibly by regulating the level of lateralization in a dose-dependent manner.

**Figure 6.**
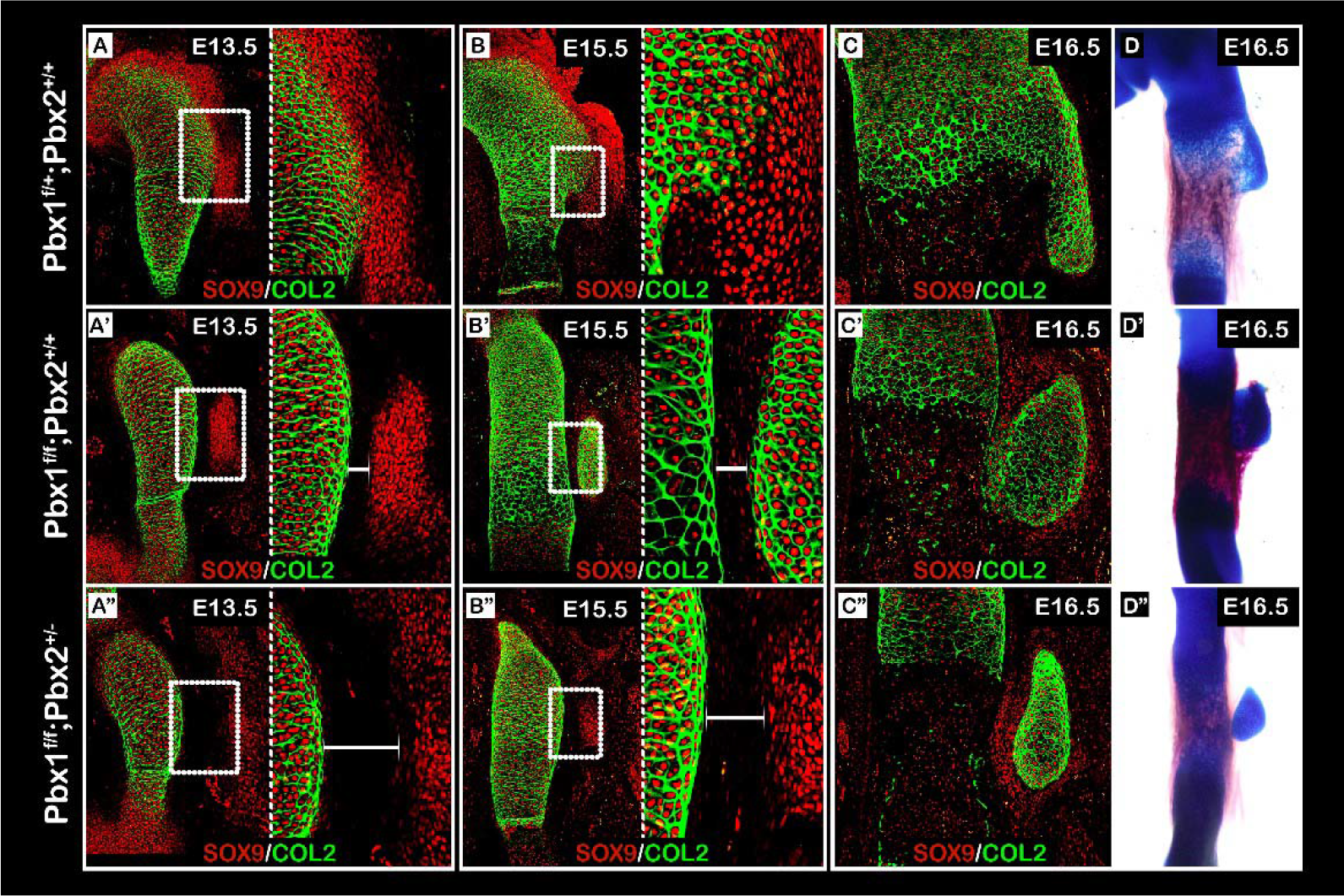
*Pbx1* and *Pbx2* coordinately regulate the patterning of proximal superstructures. **(A-C”)** Sagittal sections through the proximal humeri of E13.5 (A-A”), E15.5 (B-B”), and E16.5 (C-C”) limbs from control (A-C), *Prx1-Cre;Pbx1*^*floxed*^ (B-B”) or *Prx1-Cre;Pbx1*^*floxed*^*;Pbx2*^*het*^ (C-C”) mutant embryos that were stained against SOX9 and COL2A1. In panels A-B”, the right side of the panel is an enlargement of the humerus-DT boundary region demarcated by a white rectangle on the left. At E13.5, spatial organization of DT precursors in both mutants was abnormal (A’and A”) as DT precursors were scattered laterally away from the humeral shaft. The gap between the humeral shaft and DT precursors was twice as large in *Prx1-Cre;Pbx1*^*floxed*^*;Pbx2*^*het*^ mutants than in *Prx1-Cre;Pbx1*^*floxed*^ mutants (A’ and A”; white bars). At E15.5, whereas cells at the humerus-DT boundary of *Prx1-Cre;Pbx1*^*aoxed*^ mutants began to express high levels of *Sox9*, such expression was not observed in *Prx1-Cre;Pbx1*^*floxed*^*;Pbx2*^*het*^ mutants. Moreover, the difference in boundary region size remained consistent (B and B’; white bars). At E16.5, DT morphology ranged from attached but aplastic DT in *Prx1-Cre;Pbx1*^*floxed*^ mutants (C’) to a detached DT in *Prx1-Cre;Pbx1*^*floxed*^*;Pbx2*^*het*^ mutants (C”). **(D-D’’)** Skeletal preparations of E16.5 forelimbs further validate these diverging morphologies.

### Interaction between global and regional genetic programs fine-tune patterning of superstructures

The finding that superstructure patterning can be regulated both globally and regionally led us to hypothesize that these two programs interact with one another. To examine this possibility, we produced a compound mutant mouse carrying limb-specific knockout of *Pbx1* and *Gli3*. Specifically, on the background of a limb-specific knockout of *Pbx1* (*Prx1-Cre*;*Pbx1*^*fl/fl*^) we ablated either one or two alleles of *Gli3* (*Prx1-Cre*;*Pbx1*^*fl/fl*^; *Gli3*^*fl*/+^ or *Prx1-Cre*;*Pbx1*^*fl/fl*^;*Gli3*^*fl/fl*^, respectively). Then, sections from E13.5 and E16.5 control and mutant forelimbs were immunostained using antibodies against SOX9 and COL2A1 (Fig. 7A-B”). At E13.5, deltoid tuberosity precursors in the *Prx1-Cre*;*Pbx1*^*fl/fl*^;*Gli3f*^*fl/fl*^ embryos were laterally scattered and noticeably less condensed than in *Prx1-Cre*;*Pbx1*^*fl/fl*^;*Gli3*^*fl*/+^ embryos (Fig. 7A-A”). The phenotypic difference between genotypes became more pronounced by E16.5. Whereas ablation of a single *Gli3* allele resulted in a dysplastic deltoid tuberosity that was attached to the humeral shaft, ablation of both alleles resulted in a detached deltoid tuberosity (Fig. 7B-B”). These diverging morphologies were further validated by skeletal preparation of limbs from each mutants (Fig 7. C-C”). Collectively, these results suggest that similarly to compound *Pbx1/Pbx2* mutation, genetic interactions also occurred between *Pbx1* and *Gli3* in a dose-dependent manner, highlighting the possibility of interactions between the global and regional regulatory programs.

**Figure 7.**
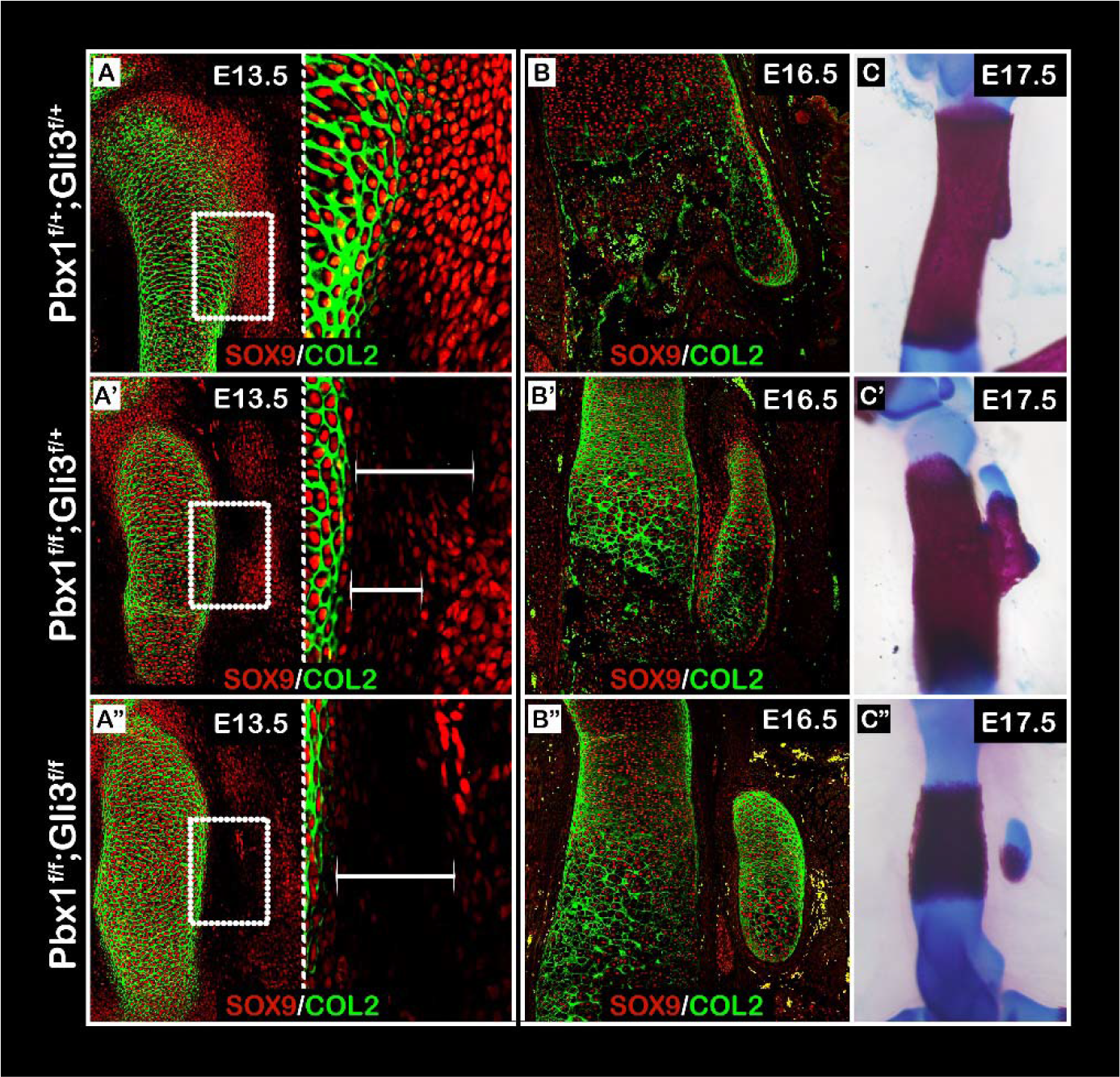
Coordinated regulation by *Pbx1* and *Gli3* highlights interaction between global and regional genetic programs. **(A-B”)** Sagittal sections through the proximal humeri of E13.5 (A-A”) and E16.5 (B-B”) limbs from control (A and B), *Prx1-Cre*;*Pbx*^*floxed*^;*Gli3*^*fl*/+^ (A’ and B’) or *Prx1-Cre*;*Pbx*^*floxed*^;*Gli3*^*floxed*^ (A” and B”) mutant embryos that were stained against SOX9 and COL2A1. (C-C”) Skeletal preparations of E17.5 forelimbs from control (C), *Prx1-Cre*;*Pbx1*^*floxed*^;*Gli3*^*fl*/+^ (C’) or *Prx1-Cre*;*Pbx1*^*floxed*^;*Gli3*^*floxed*^ (C”) mutant embryos. In panels A-A” the right side of the panel is an enlargement of the humerus-DT boundary region demarcated by a white rectangle on the left. At E13.5, spatial organization of DT precursors in both mutants was abnormal (A’,A”) as DT precursors were scattered laterally away from the humeral shaft. The gap between the humeral shaft and DT precursors was twice as large in *Prx1*-*Cre*;*Pbx1*^*floxed*^; *Gli3*^*floxed*^ mutants than in *Prx1-Cre*;*Pbx1*^*floxed*^; *Gli3*^*fl*/+^ mutants (A’,A”; white bars). At E16.5, whereas cells at the humerus-DT boundary of *Prx1-Cre*;*Pbx1*^*floxed*^; *Gli3*^*fl*/+^ mutants began to express high levels of *Sox9*, such expression was not observed in *Prx1-Cre*;*Pbx1*^*floxed*^;*Gli3*^*floxed*^ mutants. Moreover, the difference in boundary region size remained consistent (B’ and B”; white bars). At E17.5, DT morphology ranged from attached but aplastic DT in *Prx1-Cre*;*Pbx1*^*floxed*^;*Gli3*^*fl*/+^ mutants (C’) to a detached DT in *Cre*;*Pbx*^*floxed*^; *Gli3*^*floxed*^ mutants (C”).

**Figure 8.**
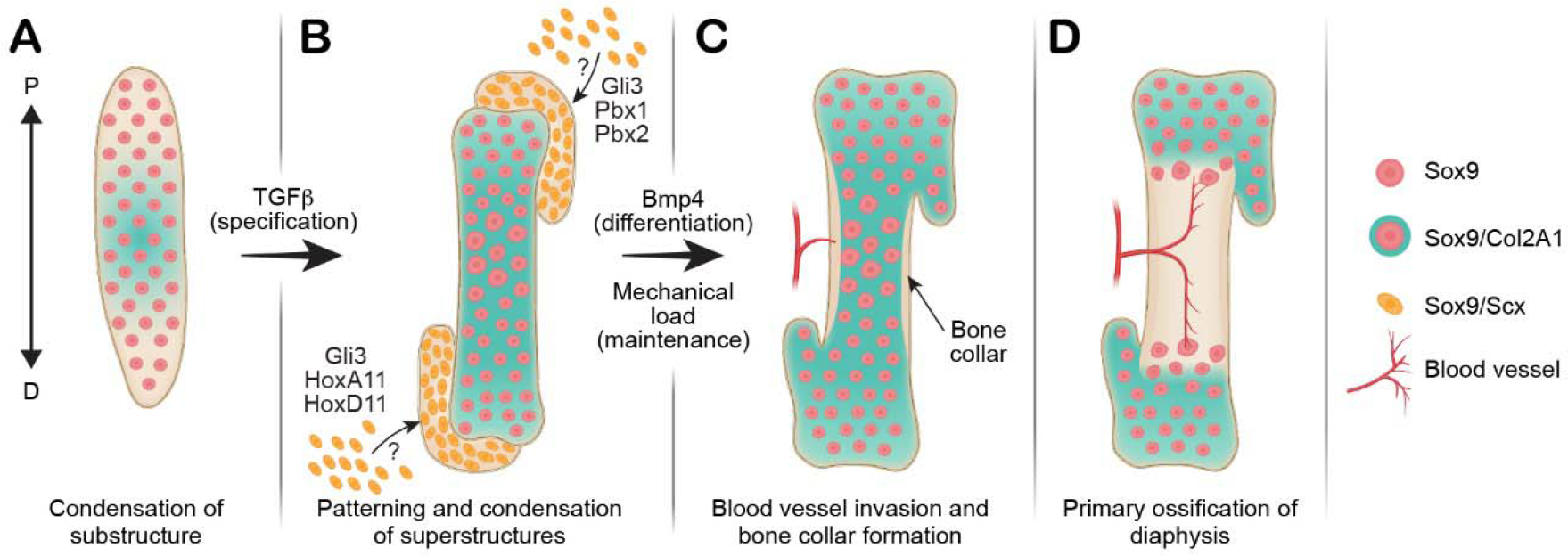
A modular model for long bone development and superstructure patterning. **(A)** Superstructure initiation is preceded by formation of the cartilaginous substructures derived from primary *Sox9*^+^ progenitors. **(B)** Subsequent to substructure differentiation, a secondary wave of *de novo* specification produces *Sox9*^+^/*Scx*^+^ superstructure progenitors. This second specification wave occurs at juxtaposition to the substructure at different time points during embryogenesis and is regulated by the TGFβ signaling pathway. The initial spatial patterning of superstructure progenitors is regulated by both global and regional molecular players, such as *Gli3* or *Pbx* and *Hox* genes, respectively. **(C)** Following specification, patterning and condensation of the superstructures precursors, they differentiate and become integral to the substructure anlagen, thus producing a complex three-dimensional cartilaginous template that is unique to each long bone. Differentiation of the superstructure is regulated by intrinsic BMP4 signaling and extrinsic mechanical stimuli. **(D)** Finally, ossification of the substructure and later of the superstructures will give rise to the morphology of the mature long bone.

## DISCUSSION

The musculoskeletal system acts as a system of levers and pulleys to create locomotion. Thus, one way to achieve variation in locomotor strategies is by changing the location of the connection between lever and pulley. Shifting the position of a bone superstructure, to which a muscle is connected by a tendon, along the bone shaft modifies the pulling force vector of that given muscle and, thereby, facilitate different type of locomotive capabilities. Variations in positioning of superstructures and their effect on locomotion have well been documented (Archer et al., 2011; Milne and O’Higgins, 2012; Polly, 2007; Salton and Sargis, 2008; Salton and Sargis, 2009). However, the mechanism that produces these variations was unknown.

An early step in the development of endochondral bones is mesenchymal condensation and the formation of the cartilaginous template that prefigures the future bone. Here, we identify components of the genetic program that regulates the subsequent step of this process, namely the patterning and condensation of superstructure precursors. Moreover, we demonstrate that the mechanism of superstructure patterning involves both global and regional modules, highlighting the modularity in long bone development. Specifically, by combining comparative transcriptomic analysis with cross-reference to existing literature, we identified a list of candidate genes that might be involved in this genetic program. Further analysis of several of these candidate genes, namely, *Gli3*, *Pbx1*, *Pbx2*, *Hoxa11* and *Hoxd11*, established their involvement in regulating superstructures patterning. Importantly, *Gli3* perturbation affected superstructures throughout different parts of the limb skeleton, whereas mutations in *Pbx1* and *Pbx2* or in *Hoxa11* and *Hoxd11* affected either proximal or distal superstructures, respectively. Interestingly, we found that the global and regional components of the genetic program are interconnected, as compound mutations of proximal regulator *Pbx1* and global regulator *Gli3* led to increment of phenotype severity, as compared to mutations in either genes alone.

In addition to these genes, our transcriptomic analysis identified 12 more candidate genes that were reportedly involved in long bone development. Interestingly, many of these genes regulate either proximal or distal limb segments, or alternatively, establish the proximodistal axis itself. For example, we identified additional HOX genes, such as *Hoxa9*, *Hoxd9*, *Hoxa10*, and *Hoxd10*, which were shown to regulate the patterning of proximal superstructures, such as the deltoid tuberosity (Wellik and Capecchi, 2003; Xu and Wellik, 2011). Other candidate genes, such as *Shox2* or *Meis1*, operate locally either as downstream effectors of HOX genes (Neufeld et al., 2014) or as cofactors of proximal regulators, such as *Pbx1* (Mercader et al., 2009), respectively. Alternatively, we identified genes that control the formation of the proximodistal axis, such as *Cyp26B1*, which modulates the levels of retinoic acid (Yashiro et al., 2004) or *Irx3, Irx5* and *Hand2*, which modulate the activity of the *Shh/Gli3* signaling pathway (Galli et al., 2010; Li et al., 2014). Together, these reports further support the modular model for superstructure patterning and suggest the involvement of at least two hierarchical programs in this process. Higher is the program that governs the establishment of the proximodistal axis and, further downstream, local programs adjust the development of each individual segment. Finally, whereas our transcriptomic analysis was designed to highlight differentially expressed genes along the proximodistal axis, it lacked information regarding globally expressed genes, which could potentially regulate superstructure patterning, such as *Gli3*, which was expressed ubiquitously in both proximal and distal *Sox9*^+^/*Scx*^+^ progenitors. Additional evidence for the existence of other global regulatory genes is given by a recent report demonstrating that *Tbx3* globally regulated patterning of forelimb superstructures, namely the greater and deltoid tuberosity and olecranon (Colasanto et al., 2016).

While our work sheds light on the genetic mechanism that regulates superstructure patterning, the cellular mechanism underlying the patterning process is still missing. A clue for its nature is given by the observations that in *Gli3*, *Pbx*, and *Hox11* mutant embryos, superstructure precursors spread abnormally away from the developing bone substructure and display delayed differentiation. This may imply the existence of a mechanism controlling the migration of *Sox9*^+^/*Scx*^+^ progenitors to specific condensation site. Once this program is perturbed, cells migrate to the wrong site, where they form an abnormal bony element. Alternatively, it is possible that *Sox9*^+^/*Scx*^+^ chondroprogenitors are specified from selected cells already present at the designated superstructure locations. In that case, the mechanism would involve activation of a chondrogenic program in specific cell subpopulations by extrinsic or intrinsic signals at specific spatiotemporal positions. In either scenario, the mechanism that decides where superstructure development should take place involves the *Gli3, Pbx* and *Hox* genes. Interestingly, several reports have demonstrated that mispatterning of specific muscles is coupled with aberrant superstructures. For example, ablation of *Tbx3* results in abnormal patterning of stylopod musculature, which attaches to dysplastic olecranon and deltoid tuberosity (Colasanto et al., 2016), whereas in *Hox11*^*aadd*^ compound mutants abnormal patterning of zeugopod musculature is accompanied by abnormal olecranon patterning (Swinehart et al., 2013). These results strongly suggest muscle and superstructure patterning may be regulated coordinately.

To conclude, our results provide insight into the formation, patterning and development of long bone superstructures and their impact on bone morphology. Furthermore, they provide strong evidence in support of the modular model of skeletogenesis and demonstrate the level of modularity during long bone morphogenesis in terms of cellular origins, genetic regulation, morphology and skeletal assembly of superstructures, namely their ability to attach or detach from the substructure. Specifically, we show that whereas *Sox9*^+^ progenitors establish the bone substructure (Fig. 7A), the superstructures anlage are formed and assembled modularly onto the substructure by *Sox9*^+^/*Scx*^+^ progenitors (Fig. 7B). We show that these unique *Sox9*^+^/*Scx*^+^ progenitors contribute not only to bone eminences but also to various condyles and sesamoid bones and that their patterning involves both global and regional regulatory modules that include *Gli3*, *Pbx* and *Hox* genes (Fig. 7B). Following their specification and patterning, these superstructures differentiate and become integral to the developing long bones (Fig. 7C and D). Importantly, by performing a comparative transcriptomic analysis, we were able to highlight additional candidate genes that may be implicated in superstructures patterning. Further studies of these molecular players will provide better understanding of the regulatory network at play in superstructure formation and long bone morphogenesis.

## MATERIALS AND METHODS

### Animals

All experiments involving mice were approved by the Institutional Animal Care and Use Committee (IACUC) of the Weizmann Institute. For all timed pregnancies, plug date was defined as E0.5. For harvesting of embryos, timed-pregnant females were euthanized by cervical dislocation. The gravid uterus was dissected out and suspended in a bath of cold phosphate-buffered saline (PBS) and the embryos were harvested after removal of the placenta. Tail genomic DNA was used for genotyping by PCR.

*Sox9-CreER*^*T2*^ mice (Soeda et al., 2010) were received from the laboratory of Haruhiko Akiyama, Kyoto University, Kyoto, Japan. *ScxGFP* transgenic mice were obtained from Ronen Schweitzer, Shriners Hospital for Children Research Division, Portland, OR, USA.

The generation of *Col2A1-CreER*^*T2*^ (Nakamura et al., 2006), *Sox9-CreER*^*T2*^ (Soeda et al., 2010), *ScxGFP* (Pryce et al., 2007), *Gli3*^*null*^ (Johnson, 1967), *Prx1-Cre* (Logan et al., 2002), floxed-*Gli3* (Blaess et al., 2008), *Pbx1*^*null*^ (Selleri et al., 2001), floxed-*Pbx1* (Ficara et al., 2008), *Pbx2*^*null*^ (Selleri et al., 2004), *Hox-A11*^*null*^ (Boulet, 2003), *Hox-D11*^*null*^ (Davis and Capecchi, 1994) and *Rosa26-tdTomato* (Madisen et al., 2010) mice has been described previously.

To create *Gli3*, *Hox11*^*aadd*^ and *Pbx1* mutant mice, animals heterozygous for the mutations were crossed; heterozygous embryos were used as a control. To create *Prx1-Gli3* or *Prx1-Pbx1* mutant mice, floxed-*Gli3* or floxed-*Pbx1* mice were mated with *Prx1-Cre-Gli3* or *Prx1-Cre-Pbx1*, respectively. To create *Prx1-Pbx1*^*flox*^-*Pbx2*^+/−^ mutant mice, floxed-*Pbx1-Pbx2*^+/-^ mice were mated with *Prx1-Cre-Pbx1*^*flox*^ mutant mice. To create *Prx1-Pbx1-Gli3* mutant mice, floxed*-Pbx1-Gli3* mice were mated with *Prx1-Cre-Pbx1-Gli3* mutant mice. As a control, *Prx1-Cre*-negative embryos were used.

For genetic lineage analysis and FACS experiments, either *Col2A1-CreER*^*T2*^ or *Sox9-CreER*^*T2*^ mice were crossed with *Rosa26-tdTomato* reporter mice. Induction of Cre recombinase was performed at various pregnancy stages by administration of 0.03 mg/gr tamoxifen/body weight in corn oil by oral gavage (stock concentration was 5 mg/ml).

### Skeletal preparations

Cartilage and bone in whole mouse embryos were visualized after skinning, disemboweling, staining with Alcian blue and Alizarin red S (Sigma) and clarification of soft tissue with 3% potassium hydroxide and 100% glycerol (McLeod, 1980).

### Paraffin sections

For preparation of paraffin sections, embryos were harvested at various ages, dissected and fixed in 4% paraformaldehyde (PFA)/PBS at 4°C overnight. After fixation, tissues were dehydrated to 100% ethanol and embedded in paraffin. The embedded tissues were sectioned at a thickness of 7 μm and mounted onto slides.

### OCT-embedded sections

For preparation of OCT-embedded sections, embryos were harvested at various ages, dissected and fixed in 1% paraformaldehyde (PFA)/PBS at 4°C overnight. Fixed embryos were then dehydrated in 30% sucrose overnight at 4°C. Next, samples were dissected, soaked in OCT for 30-60 minutes and then embedded in OCT. Frozen samples were sectioned at a thickness of 10 μm and mounted onto slides.

### Fluorescent in situ hybridization

Fluorescent in situ hybridizations (FISH) on paraffin sections were performed using digoxigenin-(DIG) labeled probes (Shwartz and Zelzer, 2014). All probes are available upon request. Detection of probes was done by anti-DIG-POD (Roche, 11207733910, 1:300), followed by Cy3-tyramide labeled fluorescent dyes, according to the instructions of the TSA Plus Fluorescent Systems Kit (Perkin Elmer). Finally, slides were counter-stained using DAPI.

### Immunofluorescence staining

For immunofluorescence staining for SOX9 and COL2A1, 7 μm-thick paraffin sections of embryo limbs were deparaffinized and rehydrated to water. Antigen was then retrieved in 10 mM citrate buffer (pH 6.0), boiled and cooked for 10 minutes in a microwave oven. In order to block non-specific binding of immunoglobulin, sections were incubated with 7% goat serum, 1% BSA dissolved in PBST. Following blockage, sections were incubated overnight at 4°C with primary anti-SOX9 antibody (1:200; AB5535; Millipore). Then, sections were washed in PBST (PBS + 0.1% Triton X-100 + 0.01% sodium azide) and incubated with Cy3-conjugated secondary fluorescent antibodies (1:100; Jackson Laboratories). After staining for SOX9, slides were washed in PBST and fixed in 4% PFA at room temperature for 10 minutes. Then, slides were incubated with proteinase K (Sigma, P9290), washed and post-fixed again in 4% PFA. Next, sections were washed and incubated overnight at 4°C with primary anti-COL2A1 antibody (1:50; DSHB, University of Iowa). The next day, sections were washed in PBST and incubated with Cy2-conjugated secondary fluorescent antibodies (1:200; Jackson Laboratories). Slides were mounted with Immuno-mount aqueous-based mounting medium (Thermo). For immunofluorescence staining for PBX1 and COL2A1, paraffin sections, antigen retrieval and blockage of non-specific binding were performed as described above. Staining for PBX1 was performed using primary anti-PBX1 antibodies (Cell Signaling, C-4342, 1:400) followed by HRP conjugated secondary antibodies (Jackson, 1:1000) and Cy3-tyramide labeled fluorescent dyes, according to the instructions of the TSA Plus Fluorescent Systems Kit. Subsequently, staining against COL2A1 was performed as described above.

For immunofluorescence staining for SOX9 and *ScxGFP*, 10 μm-thick cryostat sections of embryo limbs endogenously labeled for *ScxGFP* were used. SOX9 immunofluorescence staining was performed as described above, short of the antigen retrieval step, using primary SOX9 antibody and secondary Cy3-fluorescent antibodies.

### Tissue clearing for light sheet fluorescence microscopy

For whole-mount imaging, samples were first cleared and immunostained using the PACT-decal technique (Treweek et al., 2015; Yang et al., 2014). Briefly, either *Col2A1-CreER*^*T2*^- or *Sox9-CreER*^*T2*^-*tdTomato* mice were crossed with *Rosa26-tdTomato* reporter mice. Following tamoxifen administration at E11.5 (*Col2A1-CreER*^*T2*^) or E10.5 (*Sox9-CreER*^*T2*^), as described above, embryos were harvested at E15.5 or E14.5, respectively. Embryos were dissected and fixed in 1% paraformaldehyde (PFA)/PBS at 4°C overnight. Next, samples were washed in PBST at room temperature, then embedded into a hydrogel of 4% (wt/vol) acrylamide in 1x PBS with 0.25% thermal initiator 2,2’-azobis[2-(2-imidazolin-2-yl)propane] dihydrochloride (Wako, cat. No. VA-044). The hydrogel was allowed to polymerize at 37°C for 5 hours. The samples were removed from the hydrogel, washed in PBST, and moved to 10% SDS with 0.01% sodium azide, shaking at 37°C for 3 days, changing the SDS solution each day. Samples were washed four times with 1x PBST at room temperature over the course of a day and overnight and fixed in 4% PFA at room temperature for 10 minutes. Then, samples were incubated with proteinase K (Sigma, P9290) for 30 minutes shaking at 37°C, washed and post-fixed again in 4% PFA. To detect COL2A1, samples were first incubated with 5% goat serum dissolved in PBST overnight in order to block non-specific binding of immunoglobulin. Next, samples were incubated with primary COL2A1 antibodies (1:150) in 2.5% goat serum/PBST shaking at 37°C for 3 days, changing the antibodies solution each day. Samples were washed four times with 1x PBST at room temperature over the course of a day and overnight. Next, samples were incubated with secondary Cy3 or Cy2 antibodies (Jackson, 1:150) in 2.5% goat serum/PBST shaking at 37°C overnight. Samples were washed again with four changes of 1x PBST, and the refractive index (RI) of the sample was brought to 1.45 by submersion in a refractive index matching solution (RIMS), shaking gently at room temperature overnight. Finally, samples were embedded in 1% low gelling agarose (Sigma) in PBS in a glass capillary, submerged in RIMS and stored in the dark at room temperature until imaging.

### Light sheet fluorescence microscopy

The cleared samples were imaged with a Zeiss Lightsheet Z.1 microscope. For each limb, a low resolution image of the entire limb was taken with the 20x Clarity lens at a zoom of 0.36. Light sheet fusion of images was done if necessary (Zen software, Zeiss). Tile stitching and 3D-image reconstruction was performed using Arivis Vision4D (Arivis) and Imaris (Bitplane) Software.

### Fluorescence-activated cell sorting (FACS)

Flow cytometry analysis and sorting were performed on a BD FACS AriaIII instrument (BD Immunocytometry Systems) equipped with a 488, 407, 561 and 633 nm lasers, using a 70 μm nozzle, controlled by BD FACS Diva software v8.0.1 (BD Biosciences), at the Weizmann Institute of Science Flow Cytometry Core Facility. Further analysis was performed using FlowJo software v10.2 (Tree Star).

For collection of cells, *Sox9-CreER*^*T2*^-*tdTomato*;*ScxGFP* mice were crossed with *Rosa26-tdTomato;ScxGFP* reporter mice. Embryos were harvested at E13.5 following tamoxifen administration at E12.0, as described above. Forelimbs were dissected and suspended in cold PBS using 15 ml tubes.

To extract cells from tissues, PBS was replaced with 1 ml heated 0.05% trypsin (0.25% dissolved in DMEM, 37°C) and incubated for 15 min at 37°C, gently agitated every 5 minutes. Tissues were then dissociated by vigorous pipetting using 1 ml tips. Next, 4 ml of DMEM supplemented with 10% FBS and 1% Pen Strep was added and cell suspensions were filtered into a 15 ml tube using a syringe and a 40 μm filter net. Finally, tubes were centrifuged at 1,000 rpm for 7 minutes, supernatant was removed and cells were resuspended in 1 ml of cold PBS and used immediately for FACS.

Single-stained GFP and tdTomato control cells were used for configuration and determining gate boundaries. Live cells were gated by size and granularity using FSC-A versus SSC-A and according to staining with propidium iodide (PI, 1 μg/ml) and DAPI (1 μg/ml). FSC-W versus FSC-A was used to further distinguish single cells. In addition, unstained, GFP-stained only and tdTomato-stained only cells were mixed in various combinations to verify that the analysis excluded false-positive doublets.

### MARS-Seq

Purified RNA from FACS-isolated *Sox9*^+^/*Scx*^+^ cell samples was used for library preparation according to MARS-Seq protocol (Jaitin et al., 2014). Library quality was analyzed by 2200 TapeStation instrument (Agilent Technologies, data not shown). The experiment produced libraries of high quality and sufficient quantity. Libraries were subsequently sequenced by Illumina HiSeq 2500.

### Bioinformatic analysis

Unique molecular identifier (UMI) sequence present in the R2 read was inserted in the read name of R1 sequence file using a python script. Cutadapt was used to trim low quality and poly A/T and adapter sequences (-aAGATCGGAAGAGCACACGTCTGAACTCCAGTCAC -a "A{100}" -a "T{100}" --times 2 -q 20 -m 30). Sequences were mapped using tophat2 to mouse genome build mm10. UMI information was integrated into the BAM files as tags, using a python script. The BAM file was converted to SAM format using Samtools. Duplicate reads were marked based on having the same UMI and mapping to the same gene, using a python script. Read counts per gene were calculated using HTSeq-count and a Refseq gtf file (downloaded from igenomes UCSC), which was modified to contain a window surrounding the 3’ UTR.

DESeq2 was used for normalization and to detect differentially expressed genes based on negative binomial distribution and a generalized linear regression model. The model used contained two factors: olecranon or deltoid tuberosity *Sox9*^+^/*Scx*^+^ precursors. Genes were considered differentially expressed if the difference in expression between at least two sample types was statistically significant after adjustment with the fdrtool package (adjusted *p*-value≤0.05) (Strimmer, 2008). After filtering based on expression levels (sum normalized count in all samples greater than 10 and maximum expression level higher than 5), clustering of the standardized normalized counts was done using click algorithm (Expander package, (Ulitsky et al., 2010)). Further analysis was performed using GSEA (Broad institute) and Ingenuity (IPA).

## AUTHOR CONTRIBUTIONS

S.E. designed and carried out the experiments, analyzed the data and wrote the manuscript; S.Ku. performed FACS experiments; S.R. performed light sheet experiments; S.Kr jointly carried out experiments; K.M.P. and D.M.W. provided the *Hoxa11* and *Hoxd11* compound heterozygous mice and performed *Hoxd11 in situ* hybridization experiments; E.Z. jointly designed the study, supervised the project, analyzed and interpreted the data, and wrote the manuscript.

## ACKNOWLEDGMENTS

We thank N. Konstantin for expert editorial assistance, and members of the Zelzer laboratory for their advice and suggestions. We thank R. Schweitzer for providing the *ScxGFP* mice, and H. Akiyama for providing the *Sox9-CreER*^*T2*^ mice. We thank D. Leshkowitz, T.M. Salame, Y. addadi and O. Golani from the Weizmann Institute Department of Life Sciences Core Facilities for their guidance and assistance on conducting and analyzing the experiments and interpretation of the data. Light sheet imaging was made possible thanks to the De Picciotto-Lesser Cell Observatory in memory of Wolf and Ruth Lesser of the MICC. We thank C. Vega from the Weizmann Institute Department of Design, Photography and Printing for designing the model.

This study was supported by grants from the National Institutes of Health (NIH, #R01 AR055580), European Research Council (ERC, #310098), the Jeanne and Joseph Nissim Foundation for Life Sciences Research, the Y. Leon Benoziyo Institute for Molecular Medicine, Beth Rom-Rymer, the Estate of David Levinson, the Jaffe Bernard and Audrey Foundation, Georges Lustgarten Cancer Research Fund, the David and Fela Shapell Family Center for Genetic Disorders, the David and Fela Shapell Family Foundation INCPM Fund for Preclinical Studies, and the Estate of Bernard Bishin for the WIS-Clalit Program.

